# Age-associated B cell infiltration in salivary glands represents a hallmark of Sjögren’s-like disease in aging mice

**DOI:** 10.1101/2024.02.13.580185

**Authors:** Harini Bagavant, Justyna Durslewicz, Marcelina Pyclik, Magdalena Makuch, Joanna A. Papinska, Umesh S. Deshmukh

## Abstract

Sjögren’s disease (SjD), characterized by circulating autoantibodies and exocrine gland inflammation, is typically diagnosed in women over 50 years of age. However, the contribution of age to SjD pathogenesis is unclear. C57BL/6 female mice at different ages were studied to investigate how aging influences the dynamics of salivary gland inflammation. Salivary glands were characterized for immune cell infiltration, inflammatory gene expression, oxidative stress, and saliva production. At 8 months, gene expression of several chemokines involved in immune cell trafficking was significantly elevated. At this age, Age-associated B cells (ABCs), a unique subset of B cells expressing the myeloid markers CD11b and/or CD11c, were preferentially enriched in the salivary glands compared to other organs like the spleen or liver. The salivary gland ABCs increased with age and positively correlated with increased CD4 T follicular helper cells. By 14 months, lymphocytic foci of well-organized T and B cells spontaneously developed in the salivary glands. In addition, the mice progressively developed high titers of serum autoantibodies. A subset of aged mice developed salivary gland dysfunction mimicking SjD patients. Our data demonstrates that aging is a significant confounding factor for SjD. Thus, aged female C57BL/6 mice are more appropriate and a valuable preclinical model for investigating SjD pathogenesis and novel therapeutic interventions.

## Introduction

Sjögren’s disease (SjD) is a chronic autoimmune disorder targeting multiple organ systems [1]. The disease has a strong gender bias, predominantly affecting women over men at a 9:1 ratio [2]. The most common symptom is oral and ocular dryness caused by reduced saliva and tear production. Oral dryness in SjD is linked to difficulty in speech and swallowing food, microbial overgrowth leading to bad breath, dental decay, and tooth loss [3]. Dry eyes in SjD patients lead to a constant sensation of particulate material in the eyes, corneal damage, pain, and, in severe cases, vision loss [4]. The central nervous system involvement drives debilitating fatigue, brain fog, and depression [5]. In rare cases (<5%), patients develop life-threatening lymphomas [6]. Overall, the disease is responsible for reducing the quality of life and presents itself as a significant economic burden for the patients and their families [7]. Although genetics influence SjD susceptibility, other factors like epigenetics, sex, and environmental interactions play significant roles in disease pathogenesis [8].

SjD is more common at older ages and is typically diagnosed in or after the fifth decade of life, with the median age of diagnosis being 52 years [2]. The diagnostic hallmarks of SjD are the presence of lymphocytic foci in minor salivary gland biopsies and high titer autoantibodies to Ro60/SSA, La/SSB, and Ro52/TRIM21 proteins. It is interesting to note that autoantibodies are detectable in serum several years before the appearance of clinical symptoms and disease diagnosis [9]. However, the reasons for this delay between loss of tolerance and manifestation of clinical signs are unclear. How aging influences SjD development and progression has yet to be thoroughly interrogated, and this represents a significant gap in our knowledge of the disease.

B cells play a critical role in SjD pathogenesis, generating autoantibodies found in the systemic circulation and locally in the salivary glands. B cells contribute prominently to the lymphocytic foci of salivary gland biopsies from SjD patients. The organization of B cells within these foci into germinal center-like structures has been linked to poor disease prognosis [10]. In 2011, a unique population of B cells expressing myeloid markers was shown to accumulate in the spleens of older mice and peripheral blood of older humans [11, 12], and they were designated as Age-associated B cells (ABCs). ABCs are a heterogeneous B cell subset that lack CD21 or CD23 and express myeloid markers like CD11b and/or CD11c. The activation requirements of ABCs are distinct from conventional B cells. ABCs respond poorly to B cell receptor or CD40 ligation but expand following stimulation with Toll-Like Receptor (TLR) 7 and TLR9 agonists. Some ABCs expressing the transcription factor T-bet can produce autoantibodies and are seen in lupus-prone mice and patients with autoimmunity [13]. ABC expansion occurs preferentially in females, suggestive of their pathogenic contribution to female-dominant autoimmune diseases.

In SjD patients, peripheral blood shows an elevation of autoreactive CD21^-/low^ B cells unresponsive to CD40-mediated stimulation [14]. Moreover, CD21^-/low^ B cells expressing FcRL4, CD11c, and T-bet were recovered from minor labial salivary gland biopsies from SjD patients [15]. In mice, repeated treatment with a TLR7 agonist was associated with an expansion of T-bet**+** B cells in the spleen and an accelerated onset of SjD-like disease in NOD.B10Sn-H2b mice [16]. These reports suggested a potential pathogenic role for ABCs in salivary gland manifestations in SjD. Whether the entry of ABCs into salivary glands is a cause or consequence of lymphocytic focus formation and what mechanisms are involved in glandular infiltration are not known. To address these issues, we investigated salivary glands from aged C57BL/6 (B6) mice in the present study. Previous reports have demonstrated that aged B6 mice spontaneously develop lymphocytic infiltrates in their salivary [17, 18] and lacrimal [19] glands and autoantibodies in circulation [20]. Our study shows that early inflammatory changes in the salivary glands by 8 months of age precede lymphocytic focus formation. Further, our data demonstrate that ABC infiltration into the salivary glands inextricably links aging with loss of immune tolerance and exocrine gland inflammation.

## Methods

### Mice

Female C57BL/6N and C57BL/6J (B6) mice were obtained at different ages from the Aged Rodent Colony of the National Institute of Aging, the Jackson Laboratory (Bar Harbor, ME), and the Charles River Laboratories (Wilmington, MA). The mice were housed in a specific pathogen-free facility in the Oklahoma Medical Research Foundation vivarium (Supplementary Table S1). Mice were also bred in the OMRF facility from breeding pairs obtained from the Jackson Laboratory. All mice had unrestricted access to food (5053 PicoLab rodent Diet 20) and water. All animal experiments were approved by the Institutional Animal Care and Use Committee and followed the National Institutes of Health guidelines for the care and use of laboratory animals.

Tail bleeds were collected to determine serum autoantibody titers. Pilocarpine-induced saliva production was measured to evaluate salivary gland function [21]. Mice were euthanized at 3-, 8-9, 13-14, and 17-26 months of age. The mice were perfused with PBS before tissue collection. Submandibular salivary glands were harvested, cut into multiple pieces, and processed for gene expression analysis, histopathology, immunostaining, and flow cytometry as described previously [22, 23]. In some experiments, the spleen and liver were harvested and processed for immune cell analysis by flow cytometry.

### Salivary gland histopathology

Formalin-fixed and paraffin-embedded salivary gland sections (5 μm thickness) were stained with hematoxylin and eosin and evaluated for inflammation [24]. High-resolution images were obtained with a Zeiss AxioScan 7 digital pathology scanner, and image analysis was performed using QuPath software [25]. An aggregate of >50 lymphocytes within the glandular parenchyma was considered a focus. The sizes of lymphocytic foci and the total area of the salivary gland parenchyma were measured. The results are expressed as the area of inflammation (%) calculated as (area with lymphocytic foci/ total glandular area) X 100. A total area of 7.58 ± 0.59 sq mm (mean ±SEM) was evaluated for each sample.

### Immunofluorescence staining

Immune cell infiltrates were studied in salivary gland pieces fixed in 1% paraformaldehyde-lysine-periodate and processed as previously described [26]. The fluorochrome-conjugated monoclonal antibodies to MHC II, CD3, CD4, CD8a, and B220 were used for staining (Supplementary Table S2). For T-bet, the sections were incubated with a rabbit anti-T-bet antibody followed by a fluorochrome-conjugated donkey anti-rabbit IgG antibody. DAPI was used for nuclear staining, and the tissues were mounted in Prolong Gold (Invitrogen, Waltham, MA). Images were captured using the LSM710 confocal microscope with Zen software (Carl Zeiss Microscopy LLC, White Plains, NY).

### Gene expression analyses

Snap-frozen salivary gland pieces in liquid nitrogen were used for gene expression analysis. RNA was isolated using the RNeasy mini kit (Qiagen, Germantown, MD), and gene expression was studied using the nCounter mouse myeloid cell panel with probes for 734 genes and 20 internal reference genes (Nanostring Technologies, Seattle, WA). Gene expression analysis was performed using the ROSALIND analysis software (ROSALIND, San Diego, CA). Ingenuity Pathway Analysis (Qiagen) was also performed using differentially expressed genes.

### Flow Cytometry

The salivary glands, livers, and spleens were minced and digested with collagenase D (1mg/ml) and DNase (0.1mg/ml) (Sigma Aldrich, St Louis, MO) to obtain single-cell suspensions. Using standard protocols, cells were stained with fluorochrome-conjugated antibodies (Supplementary Table S2). Zombie Aqua fixable viability kit (Biolegend, San Diego, CA) was used for live/dead cell discrimination. Sample acquisition was performed on a 5-Laser Cytek Aurora spectral flow cytometer using Spectroflo software (Cytek Biosciences, Fremont, CA). FlowJo software (Becton Dickenson, Ashland, OR) was used for data analysis. Dimensionality reduction was performed with the Uniform Manifold Approximation and Projection (UMAP) plug-in, and the X-shift algorithm was used for clustering B cell subsets.

### Serum antibody analysis

Serum antibodies to nuclear and cytoplasmic antigens were evaluated by indirect immunofluorescence staining of HeLa cells, as described previously [24]. Serum samples were serially diluted starting at 1:50 and tested for reactivity. FITC-conjugated goat F(ab)2 anti-mouse IgG (Jackson Immunoresearch) was used to detect bound antibodies. Nuclei were stained with DAPI and mounted in ProLong Gold (Invitrogen). Images were captured at fixed exposure on a Zeiss Axiovert fluorescence microscope using AxioVision software. The results are expressed as endpoint titers (highest serum dilution generating a signal over the negative control). All experiments used a pooled serum sample from young (8-week) C57BL/6 mice at 1:50 dilution as a negative control.

### Statistical analysis

All statistical analyses were performed using Prism 9.4 software (GraphPad, San Diego, CA). Gaussian distribution was assessed for each dataset using a normality test. Student’s t-test was used to compare differences between two groups when data followed a Gaussian distribution. Two-way analysis of variance (ANOVA) followed by Sidak’s post-test was used for multiple comparisons. The non-parametric Mann-Whitney test determined differences between two groups for non-Gaussian distributions. Kruskal-Wallis test, followed by Dunn’s post-test, was used for multiple comparisons. The Spearman method was used to determine correlation coefficients. A p-value of less than 0.05 was considered statistically significant.

## Results

### Hierarchical recruitment of adaptive immune cells into aging salivary glands is preceded by changes in chemokine gene expression in B6 mice

Submandibular salivary glands harvested from 3-, 8-, 13- and >17-month-old C57BL/6N (open circles) or C57BL/6J (filled circles) female mice were evaluated for the formation of lymphocytic foci (defined as an aggregation of >50 lymphocytes), and the area of inflammation was quantified (Figure 1A). None of the mice showed lymphocytic foci at 3 months (Figure 1B, 1F). Mild inflammation was first detected at 8-9 months of age in 3/14 mice. By 13 months, the glands showed lymphocytic foci in the salivary gland parenchyma (Figure 1C-E). The foci were of varying sizes and were present in the peri-vascular and peri-ductal regions extending into the acini. The incidence of inflammation significantly increased with age, progressing from 6/10 mice at 13 months and 13/17 mice at >17 months. Immunostaining showed that the lymphocytic foci comprised T cells (CD4 and CD8) and B cells (Figure 1G). Large foci showed well-organized, discrete areas of T cells (both CD4 and CD8) and B cells (Figure 1H). Increased MHC II expression was seen in the foci, corresponding to the B cell areas (Supplementary Figure S1).

**Fig. 1.**
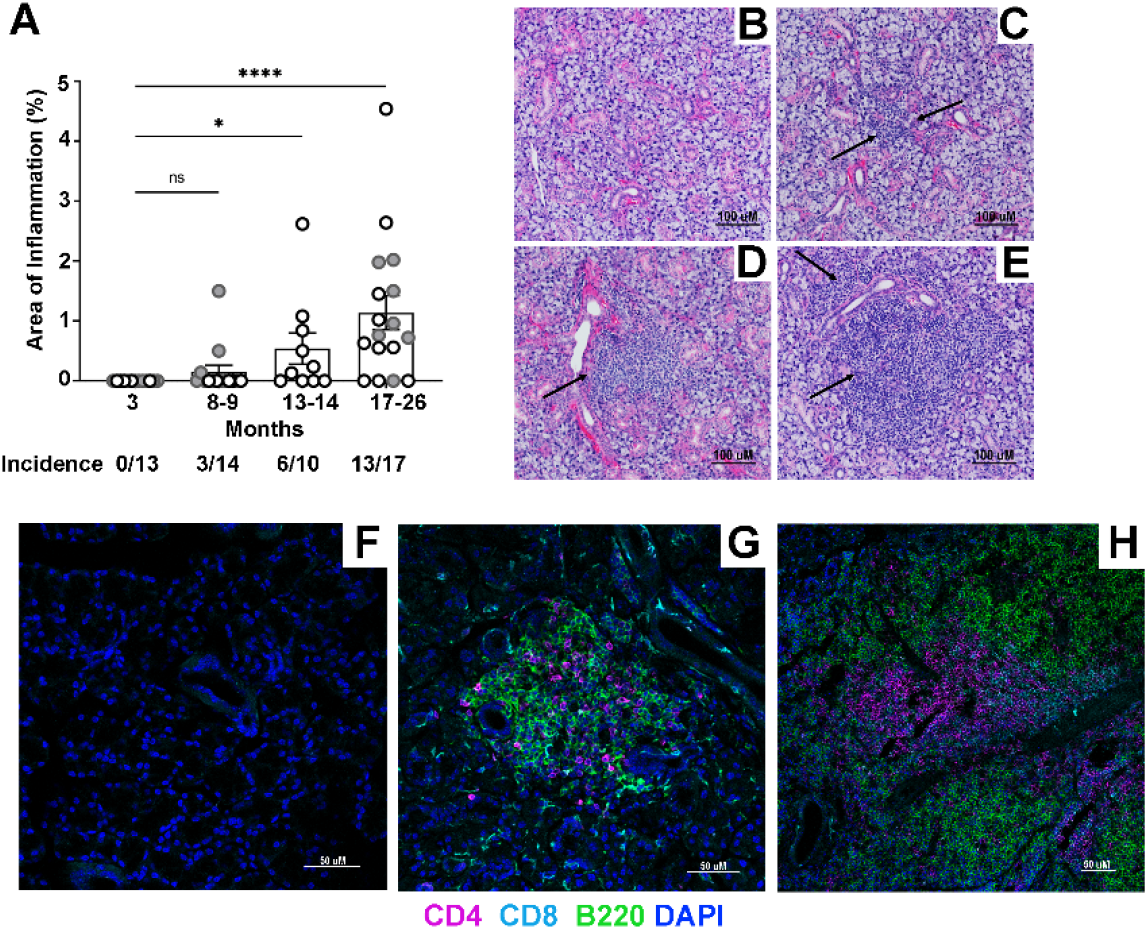
C57BL/6 mice develop salivary gland inflammation with increasing age. **A**. The area of inflammation in salivary glands from C57BL/6N (open circles) and C57BL/6J (closed circles) at different ages was calculated as (the size of lymphocytic foci/total area of the salivary gland section) x 100. Each data point represents one mouse. The difference between the groups was determined by Kruskal-Wallis one-way ANOVA and Dunn’s post-test for multiple comparisons. ns, not significant; *\* p*<0.05; \*\*\*\**p*<0.0001. **B-E**. Representative images of H&E stained sections from a normal salivary gland in 3-month-old **(B)** and inflamed glands in 13-month-old mice **(C-E)** showing peri-vascular and per-ductal lymphocytic foci (arrows) of different sizes invading the surrounding acini and ducts. Scale bar = 100 μm. **F-H**. Representative images of salivary gland sections stained with antibodies to CD4, CD8, and B220 at 3 months **(F)** and at 13 months, showing a small focus **(G)** and a large focus **(H)**. The large focus shows distinct areas of T and B cells, suggesting organization into germinal center formation. Scale bars = 50μm

The pathology of salivary gland inflammation was similar in both the B6 substrains. However, the C57BL/6N and C57BL/6J have numerous phenotypic and genotypic differences [27]. Therefore, all gene expression and flow cytometry analyses presented in this study are restricted to C57BL/6N mice, which also is the substrain provided by the National Institute of Aging, Aged Rodent Colony and is widely used in aging research.

To investigate the mechanisms for the onset of inflammation, RNA was isolated from submandibular glands of mice at 3-, 8-, and 13-months (n=4/grp), and gene expression was studied using the nCounter myeloid gene expression probe set. Hierarchical clustering and principal component analysis showed age-dependent gene expression patterns in 11/12 mice. One mouse in the 13-month age group fell between the 8- and 13-month groups (Supplementary Figure S2A and S2B). This pattern was replicated in the hierarchical clustering based on differential gene expression (DEG) (Figure 2A-C). Significant changes in DEG were noted between 3- and 8-month salivary glands, with 23/734 upregulated and 9/734 downregulated genes (Figure 2D). Despite the absence of lymphocytic foci, 8-month-old salivary glands showed increased expression of chemokines (*Cxcl9, Cxcl11, Ccl5, Ccl8*) and monocyte/macrophage-related genes (*Trem2, Ldlr, Cebpd*). At 13 months, there was a further increase in the number of genes and the extent of fold change in DEG (Figure 2E, 2F). Notably, the most significant gains occurred in chemokine expression and MHCII, suggesting immune cell-related inflammation. Thus, with aging, pro-inflammatory gene expression change drives the spontaneous recruitment of immune cells in salivary glands.

**Fig. 2.**
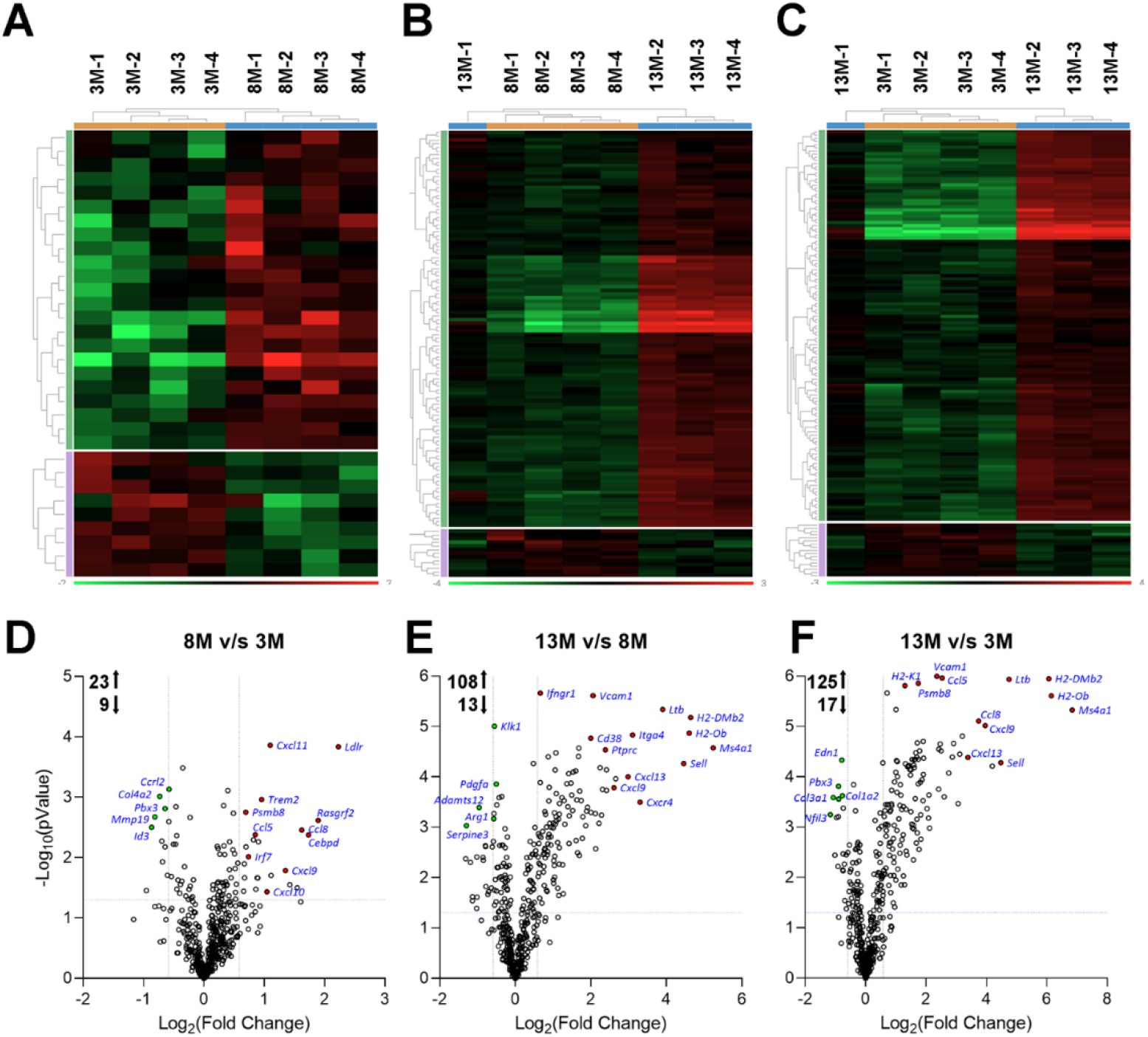
Gene expression analysis in salivary glands of mice at 3-, 8- and 13-months of age. **A-C**. Heat maps showing hierarchical clustering of DEG in salivary glands from 3-, 8-, and 13-month-old female mice, n=4/group. Each row represents a gene, and each column represents a mouse. Based on gene expression patterns, mice from each group cluster together except for one mouse at 13 months of age. **D-F**. Volcano plots showing DEG in salivary glands from mice at different ages. The DEG showing maximal fold changes and highest significance are labeled.

Commensurate with the increased chemokine expression, flow cytometry analysis of the salivary glands at 3-, 8-, 13- and > 17 months showed a progressive expansion of the adaptive immune cells, specifically in CD4 T, CD8 T, and B cell populations (Figure 3A). The UMAP multicolor plot overlays and gating strategy for salivary gland immune cell analysis are shown in Supplementary Figure S3. Compared to 3 months, 13-month-old salivary glands showed a significant increase in CD4 (3.99x) and CD8 (4.99x) T cells. The most dramatic change was in the B cells with a 33.5x increase (Figure 3B). Thus, the spontaneous salivary gland inflammation in aging B6 mice shows characteristics similar to the lymphocytic infiltration in minor labial gland biopsies from SjD patients [28].

**Fig. 3.**
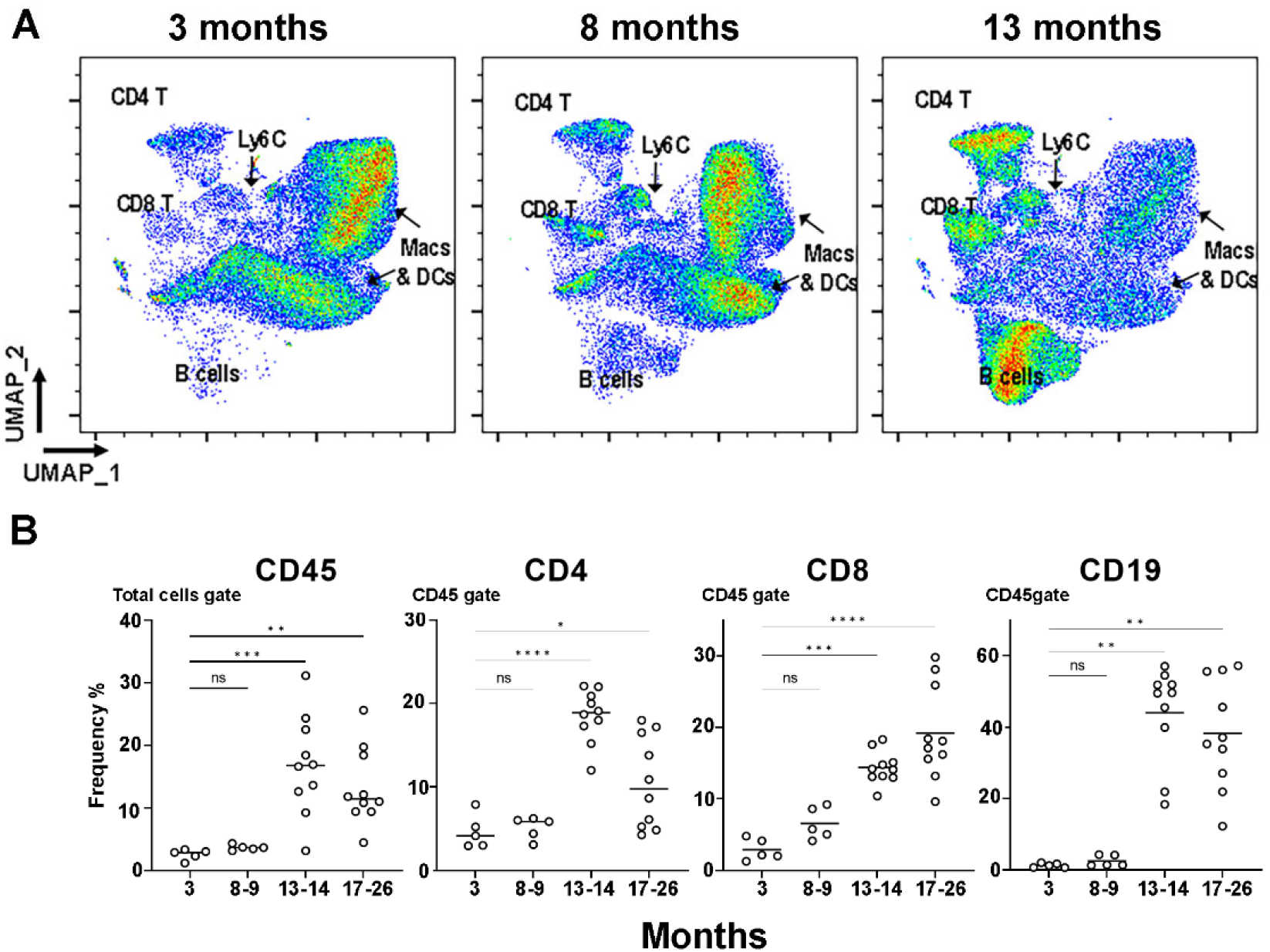
Immune cells infiltrating the salivary glands at different ages were analyzed by flow cytometry. **A**. UMAP plots identifying CD4 T, CD8 T, Ly6C monocyte, B cell, and macrophage/ dendritic cell clusters at 3-, 8-, and 13-months. Each plot represents 50,000 CD45+ cells from 5 mice with 10,000 CD45+ cells per mouse. **B**. Frequency of different immune cell types in salivary glands at 3 months (n=5), 8-9 months (n=5), 13-14 months (n=10), and 17-26 months (n=10) of age. ns, not significant; *\* p*<0.05; \**\* p*<0.01; \*\*\**p*<0.001,\*\*\*\**p*<0.0001.

### Age-associated B cells (ABCs) accumulate within lymphocytic infiltrates in aging salivary glands

CD19+B220+ B cells were analyzed for CD21, CD23, CD11b, and CD11c expression to characterize B cell subsets infiltrating the salivary glands (Figure 4A). The gating for B cell subsets in the spleen at 13 months (Supplementary Figure S4) was used to define Follicular B cells (CD23+CD21+), Marginal Zone B cells (CD23^neg^CD21^hi^), and ABCs (CD21^neg^CD23^neg^ B cells expressing CD11b/CD11c). ABCs were detected in the salivary glands at 8 months and progressively increased in frequency at 13- and >17 months of age (Figure 4B). Thus, ABCs were recruited into the salivary glands before the formation of large lymphocytic foci. Interestingly, salivary gland sections from aged mice showed that ABCs (T-bet+CD19+) were present within the lymphocytic focus (Supplementary Figure S5).

**Fig. 4.**
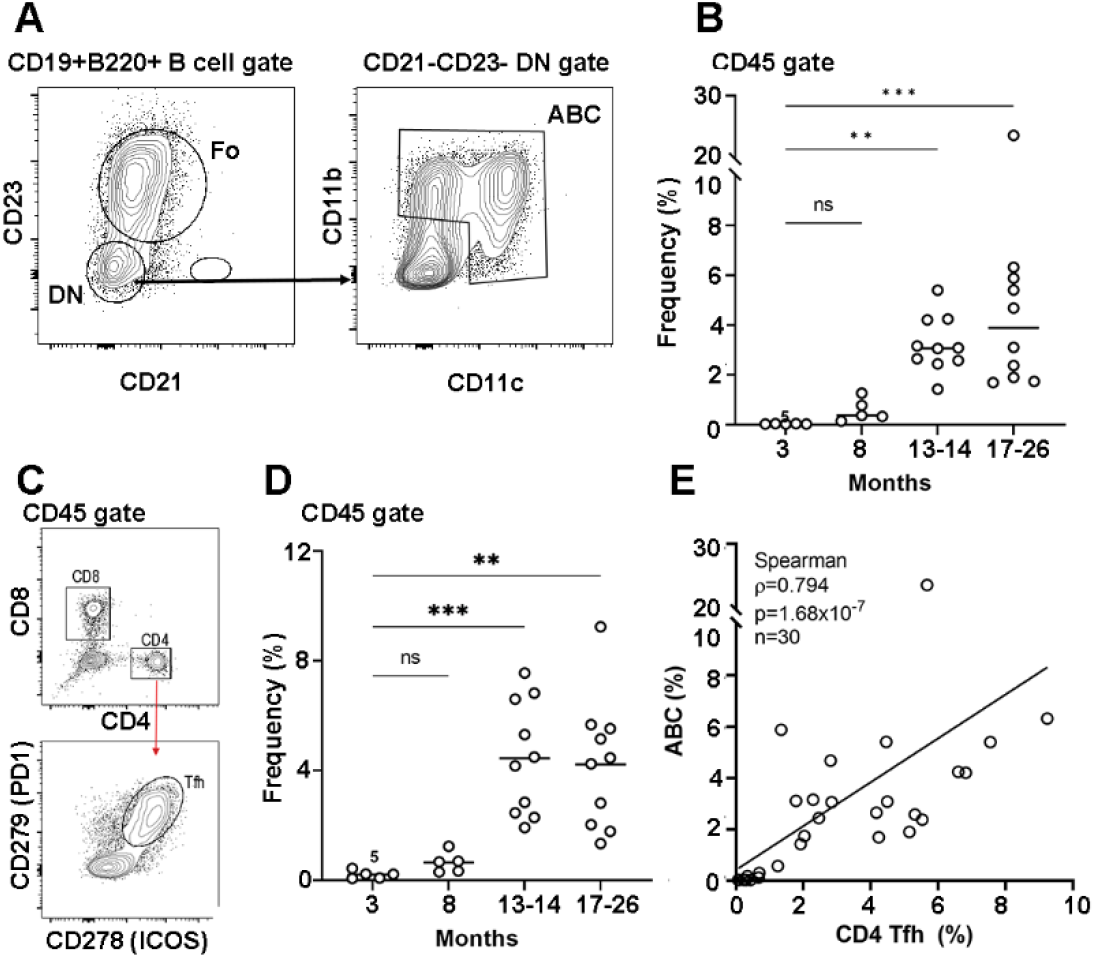
Increased Age-associated B cells (ABCs) accumulation in salivary glands of aged mice correlates with increased T follicular helper (Tfh) cells. ABCs **(A-B)** and Tfh **(C-D)** cells from salivary glands at 3 months (n=5), 8-9 months (n=5), 13-14 months (n=10), and 17-26 months (n=10) of age. **A**. Gating strategy showing CD23+CD21+ Follicular cells (Fo) and CD21^neg^CD23^neg^ Double Negative (DN) cells within the B cell gate. ABCs were identified as the DN cells expressing either CD11b and/or CD11c. **B**. Frequency of ABCs within the CD45+ gate. **C**. Gating strategy showing CD4 and CD8 T cells within the CD45+ gate. Tfh were identified as CD4 T cells expressing CD279 (PD1) and CD278 (ICOS). **D**. Frequency of Tfh within the CD45+ gate. **E**. Correlation between frequencies of ABCs and Tfh in salivary glands determined by the Spearman method. Each data point represents one mouse. In **(B)** and **(D)**, the difference between the groups was determined by Kruskal-Wallis one-way ANOVA and Dunn’s post-test for multiple comparisons. ns, not significant; \**\* p*<0.01; \*\*\**p*<0.001.

The expansion and differentiation of ABCs into antibody-producing cells may occur with or without help from T follicular helper (Tfh) cells in a germinal center-dependent or independent fashion [29]. To determine the presence of Tfh cells, CD4 cells in the salivary glands were interrogated for CD278 (ICOS) and CD279 (PD-1) expression (Figure 4C). Tfh cells were present in the glands by 8 months and increased significantly with age (Figure 4D). Indeed, a strong positive correlation existed between ABCs and CD4 Tfh frequencies (Figure 4E).

To determine whether the ABCs-CD4 Tfh correlation was indicative of a germinal center interaction, salivary gland B cells were analyzed for GL-7 expression. At 13-14 months, GL-7 positive cells constituted 1.67 ± 0.19 % (mean ± SEM, n=10) of the B cell subset. A considerable fraction of the GL-7+ germinal center B cells (26.02 ± 3.88 %) were CD21^neg^CD23^neg^CD11b+CD11c+/-. Thus, tertiary lymphoid follicle-like structures in the salivary glands consisted of a mixture of ABC and non-ABC germinal center B cells.

### The aging salivary glands provide a favorable environment for ABC recruitment and expansion

To investigate whether the ABC infiltration was unique to the salivary glands, B cells in the liver and spleen were studied at 8 and 14 months of age (Figure 5A-5C). To compare the different B cell subsets, B cells were clustered using UMAP and X-shift algorithms based on CD23, CD21, CD11c, and CD11b expression (Supplementary Figure S6). At 8 months, a significant fraction of B cells in the salivary glands were ABCs (24.3 ± 3.54) compared to the spleen (1.62 ± 0.27) or liver (1.39 ± 0.21; *p*<0.0001, n=5). At 14 months, the ABC fractions were comparable between the different organs. Thus, ABCs are preferentially recruited into the salivary gland before the formation of lymphocytic foci.

**Fig. 5.**
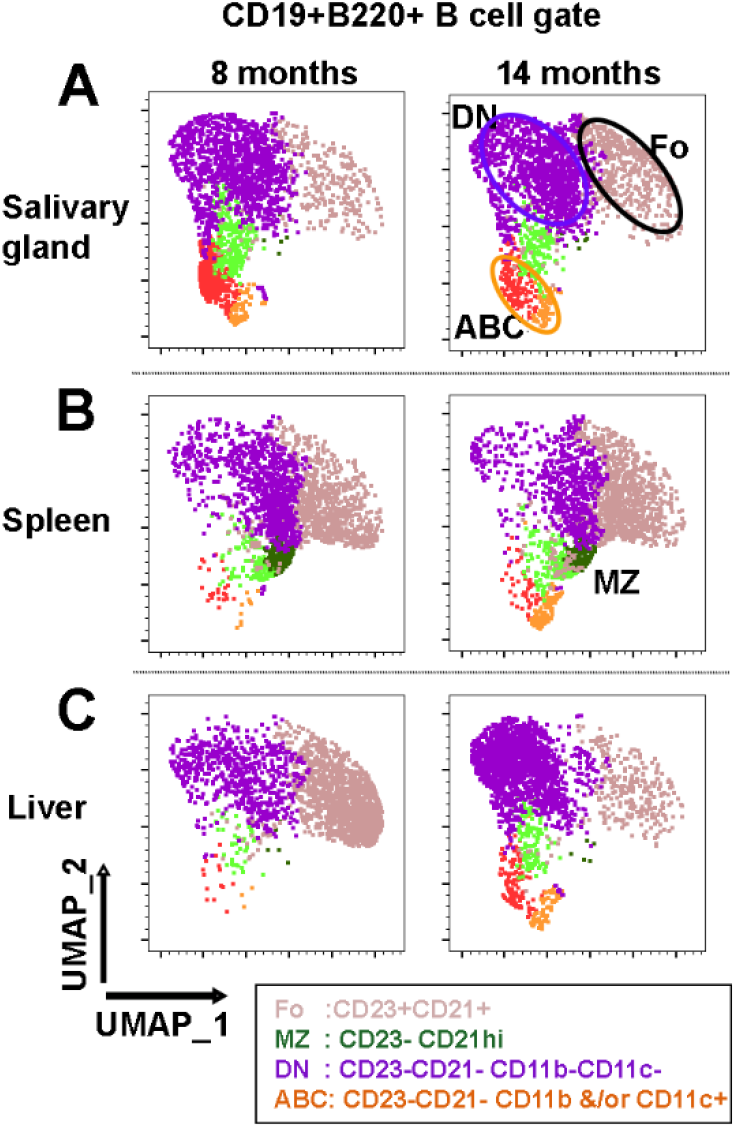
Analysis of B cell subsets at 8- and 14-months of age. **A**. Salivary glands. **B**. Spleen. **C**. Liver. UMAP plots of CD19+B220+ B cells clustering based on CD21, CD23, CD11c, and CD11b expression. Multicolor plot overlays are shown in Supplementary Figure S6. Clusters were defined by the X-Shift clustering algorithm. Each plot represents 2,500 B cells from 5 mice, with 500 B cells per mouse. CD23^neg^CD21^hi^ marginal zone (MZ) cells were seen predominantly in the spleen and rarely in other organs.

### Aged C57BL/6 mice spontaneously develop high titers of autoantibodies in serum

Loss of tolerance with spontaneous development of autoantibodies has been reported in aged B6 mice [20]. In the present study, sera collected during sacrifice were tested for antinuclear antibodies (ANA) in an indirect immunofluorescence assay (Figure 6). Aged mice showed antibodies with heterogeneous specificities to nuclear and cytoplasmic antigens (Figure 6B-D). Endpoint titers showed significant elevation in serum autoantibody with increasing age (Figure 6E). Overall, these results demonstrate that with aging, there is a loss of immunological tolerance to self-antigens in the B6 mice.

**Fig. 6.**
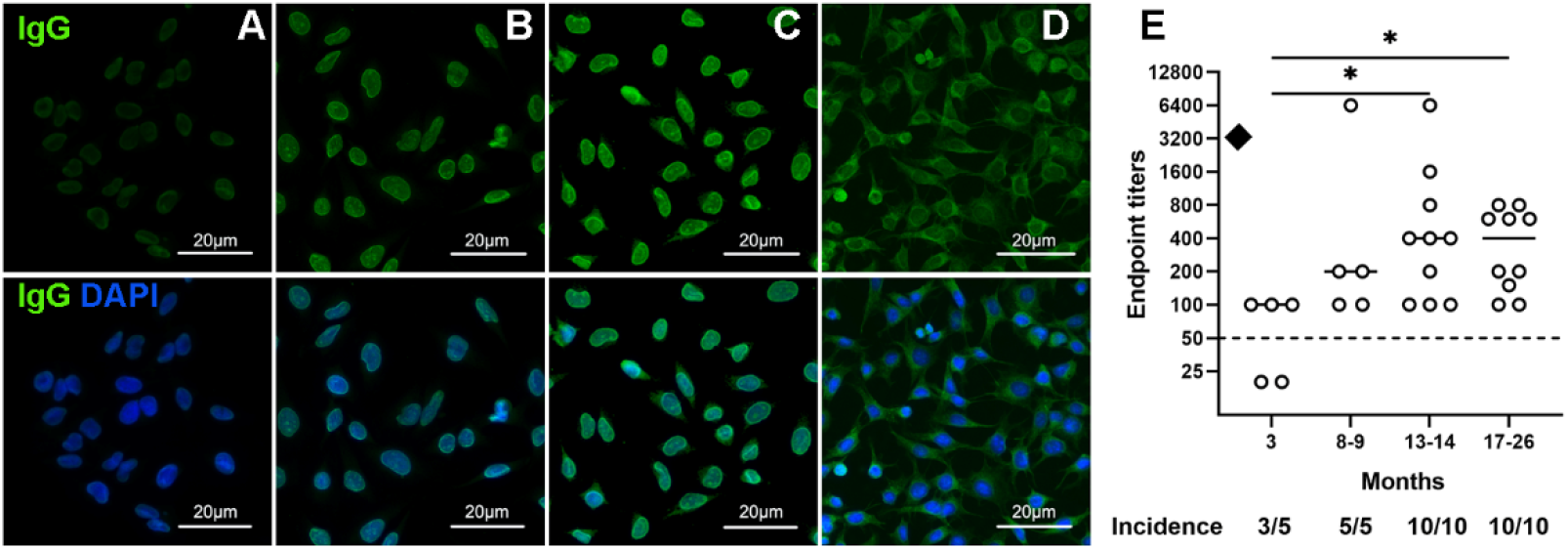
Aged C57BL/6 mice spontaneously develop autoantibodies to nuclear and cytoplasmic antigens. The top panel shows autoantibody reactivity in sera of C57BL/6 females, and the bottom panel shows overlay with nuclear stain (DAPI). **A**. Fluorescence intensity with a negative control serum at 1:50 dilution. **B-E**. Different reactivity patterns to the nuclear membrane **(B)**, homogenous nuclear **(C)**, and cytoplasmic antigens **(D)**. Scale bar= 20 microns. **E**. Endpoint titers were determined by serial dilution. ♦: represents end-point titer of pooled sera from autoimmune MRL.*lpr/lpr* mice. The difference between the groups was determined by Kruskal-Wallis one-way ANOVA and Dunn’s post-test for multiple comparisons. ns, not significant; \**p*<0.05.

## Discussion

The clinical diagnosis of Sjögren’s Disease occurs most frequently in the 5^th^-6^th^ decade of life, with symptoms worsening with age. This human life phase is equivalent to 14-18 month old mice [30]. Our results show that by 14 months, female B6 mice spontaneously develop well-organized lymphocytic foci within the salivary glands. No significant differences were noted in salivary gland inflammation between C57BL/6J and C57BL6/N mouse strains, and the kinetics of salivary gland inflammation and autoantibody development in our study were comparable with other reports in B6 mice [17, 18]. Collectively, these data establish that in B6 mice, aging of the exocrine glands facilitates adaptive immune cell infiltration.

Our study shows that by 8 months of age, corresponding to the 4^th^ decade in humans, the salivary glands from B6 mice underwent inflammatory changes. The chemokine *Cxcl11* showed significant upregulation. CXCL11 is a high-affinity ligand for CXCR3, a chemokine receptor on CD4 Th1 and CD8 effector T cells [31]. The CXCL11-CXCR3 axis is essential for recruiting activated CD4 and CD8 T cells and setting up a pro-inflammatory amplification loop for lymphocytic infiltration. The salivary glands of SjD patients have been shown to express CXCL11 [32], and CXCL11 levels were elevated in the tear film and on the ocular surface in SjD patients [33]. The study showing that disrupting CXCR3 function in NOD mice significantly influenced the development of SjD-like disease further highlights the importance of the CXCL11-CXCR3 axis in SjD [34]. The mechanisms involved in elevated CXCL11 expression in salivary glands of aging mice are unknown. Previous studies have suggested that in age-related macular degeneration, advanced glycation end-products affect CXCL11 expression in the retinal pigmental epithelium [35]. Whether advanced glycation end-products accumulate in aging salivary glands and influence CXCL11 expression by salivary gland epithelial cells will be investigated in the future.

Similar to our finding, chemokines produced by aged salivary epithelial cells have been implicated in inducing pro-inflammatory cytokines and T-cell migration [18]. In this report, the authors suggested that aging salivary glands accumulated PD1+ senescent T cells. Although, in our study, we did not specifically investigate T cell senescence, by 14 months, the gene expression patterns showed significantly increased Th1 and Th2 T cell function, classical and alternative macrophage activation, and cytokine storm signaling pathways (Supplementary Figure S7). However, our results showed that PD1-PDL1 signaling pathways associated with T-cell exhaustion and/or senescence were downregulated (Supplementary Figure S7).

To our knowledge, this is the first study demonstrating ABCs infiltrating the salivary glands of aging B6 mice. At 8 months of age, the frequency of ABCs in salivary glands was higher than that of the spleen and liver. The mechanisms responsible for the early enrichment of ABCs in salivary glands are unknown. However, given that the engagement of TLR7 plays a vital role in ABC expansion, the possibility exists that endogenous ligands in the salivary glands sustain or expand the ABC population. Salivary gland epithelial cells have been shown to secrete exosomes loaded with SjD-associated Ro-RNP antigens [36]. The small cytoplasmic RNA associated with the Ro-RNP activates TLR7 [37] and might be involved in the expansion of ABCs in the salivary gland. Additionally, our study noted a strong correlation between the frequencies of ABCs and Tfh in the salivary glands of aging mice. These data suggest that Tfh also might be involved in expanding the ABCs population in salivary glands.

How the heterogeneity within ABC populations reported in different autoimmune and infectious diseases [38, 39] affects SjD is unknown. In SjD patients, the CD21^-/low^ B cells elevated in peripheral blood showed immunoglobulin gene rearrangement indicative of memory B cells and expressed antibodies reactive with nuclear and cytoplasmic autoantigens. In minor labial gland biopsies from SjD patients, a heterogeneous population of FcRL4+ B cells expressed Ki67, indicating actively dividing cells [40]. A fraction of the FcRL4+ B cells within the peri-ductal infiltrates were CD11c+Tbet+ and bonafide ABCs [15]. Thus, the pathogenic contribution of ABCs in SjD may range from autoantibody production to lymphomagenesis. Indeed in a mouse model of SjD, ABCs isolated from spleens and activated in vitro, secreted antibodies reactive with multiple self-antigens [41].

Aging in B6 female mice results in salivary gland inflammation and systemic autoimmunity, critical characteristics of SjD. Salivary gland dysfunction, as measured by pilocarpine-induced saliva production, was reduced in 1/5 mice at 13 months and 4/10 mice at 17-26 months of age. However, salivary function did not show a statistically significant drop between the groups (Figure 7A). Further, the drop in saliva volumes did not correlate with the severity of inflammation (Figure 7B), suggesting a significant reserve in functional tissue. In contrast, tear production in aging B6 mice showed increased tear production despite severe lacrimal gland inflammation [19], suggesting differences in functional outcomes between salivary and lacrimal glands.

**Fig. 7.**
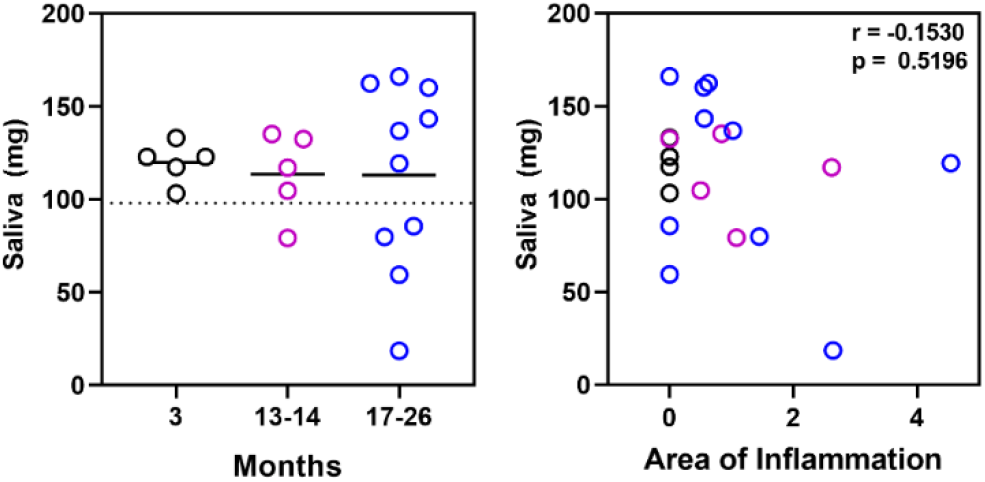
Age-associated decline in salivary gland function develops in some mice and is not proportional to the extent of inflammation. **A**. Pilocarpine-induced saliva in mice at different ages showed reduced saliva production in some older mice. The cut-off was calculated as mean-2SD of the control mice at 3 months of age. **B**. In aged mice, the functional decline did not correlate with the severity of salivary gland inflammation.

The number of mouse models that recapitulate some features of SjD is abundant in the literature [42]. However, a lack of consideration of age as a confounding factor is a significant caveat in most models. Some of the more popular models on the non-obese diabetes (NOD) genetic background develop salivary gland inflammation as early as 10-12 weeks of age [43], corresponding to a 20-30-year-old human being. Since female B6 mice develop severe age-dependent exocrine gland inflammation and some functional loss, mimicking SjD patients, aged mice would be more appropriate preclinical models for testing immunomodulating therapies.

## Acknowledgments

The authors thank the OMRF Imaging Core and Flow Cytometry Core for their assistance. In addition, we thank the Laboratory for Molecular Biology and Cytometry Research at Oklahoma University Health Sciences Center for using the Core Facility, which provided Nanostring gene expression service. Authors are thankful to the Aged Rodent Colony Resource of the National Institute of Aging for providing C57BL/6N mice at different ages.

## Author Contributions

HB designed and performed experiments, analyzed the data, and wrote the manuscript. JD, MP, MM, and JP performed experiments and analyzed data. MP, MM, and JP edited the manuscript. USD conceived the idea, designed experiments, and wrote the manuscript. All authors contributed to the article and approved the submitted version.

