## Supplementary material for "Age-associated B cell infiltration in salivary glands represents a hallmark of Sjögren’s-like disease in aging mice": Supplemetary Figures and Tables

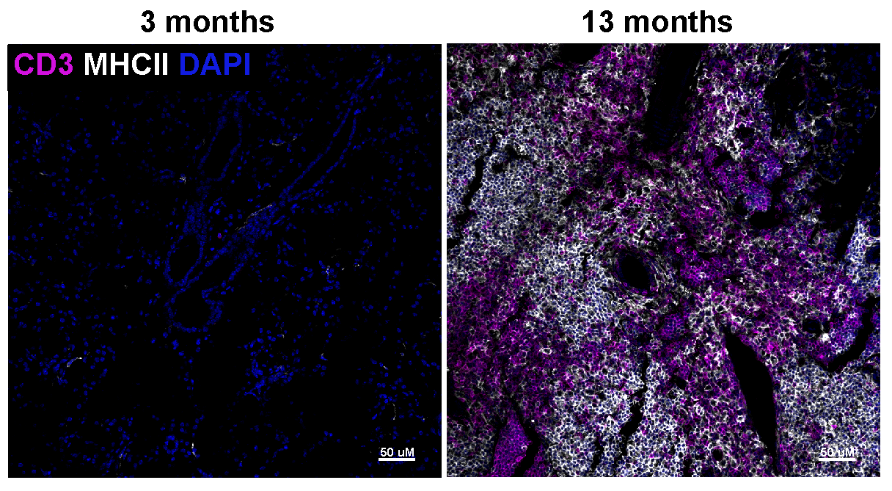

**Supplementary Figure S1.** Representative images of salivary gland sections stained with antibodies to CD3 and MHC II at 3 months and 13 months show upregulated MHCII expression in areas of B cell clustering within the lymphocytic focus. Scale bar=50 microns.

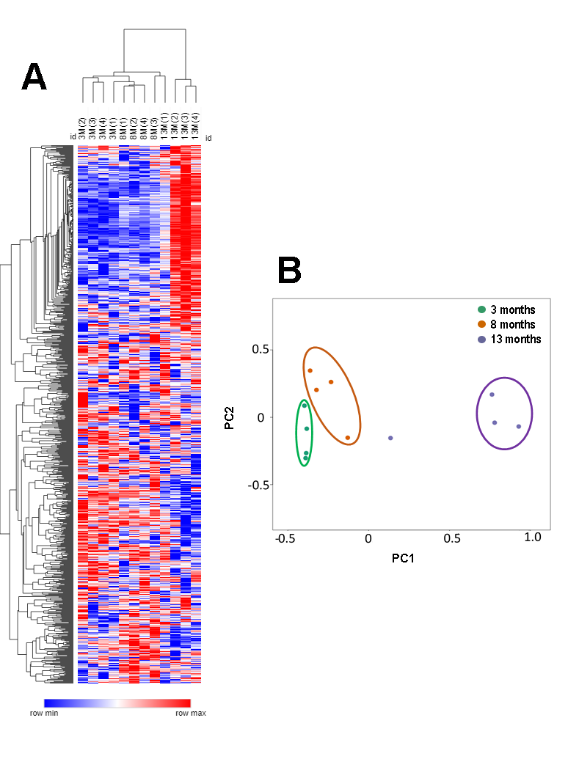

**Supplementary Figure S2. (A)** Heat Map showing hierarchical clustering of gene expression in salivary glands of mice at 3 months (3M), 8 months (8M), and 13 months (13M) of age, n=4/group. Each row represents one gene, and each column represents one mouse. **(B)** Principal component analysis shows an age-dependent clustering of gene expression patterns. Some heterogeneity is present at 13 months, where one mouse (m1) falls in between the 8- and 13-month groups.

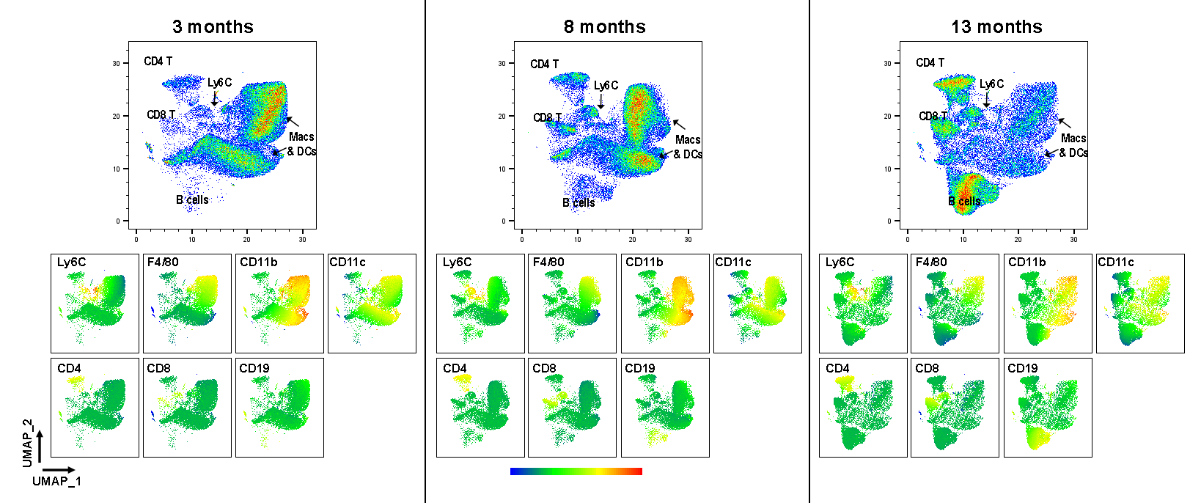

**Supplementary Figure S3A**. UMAP plots (top) with multicolor plot overlays (bottom) identifying CD4 T, CD8 T, Ly6C monocyte, B cell, and macrophage/ dendritic cell clusters at 3-, 8- 13- months. Each plot represents 50,000 CD45+ cells from 5 mice with 10,000 CD45+ cells per mouse.

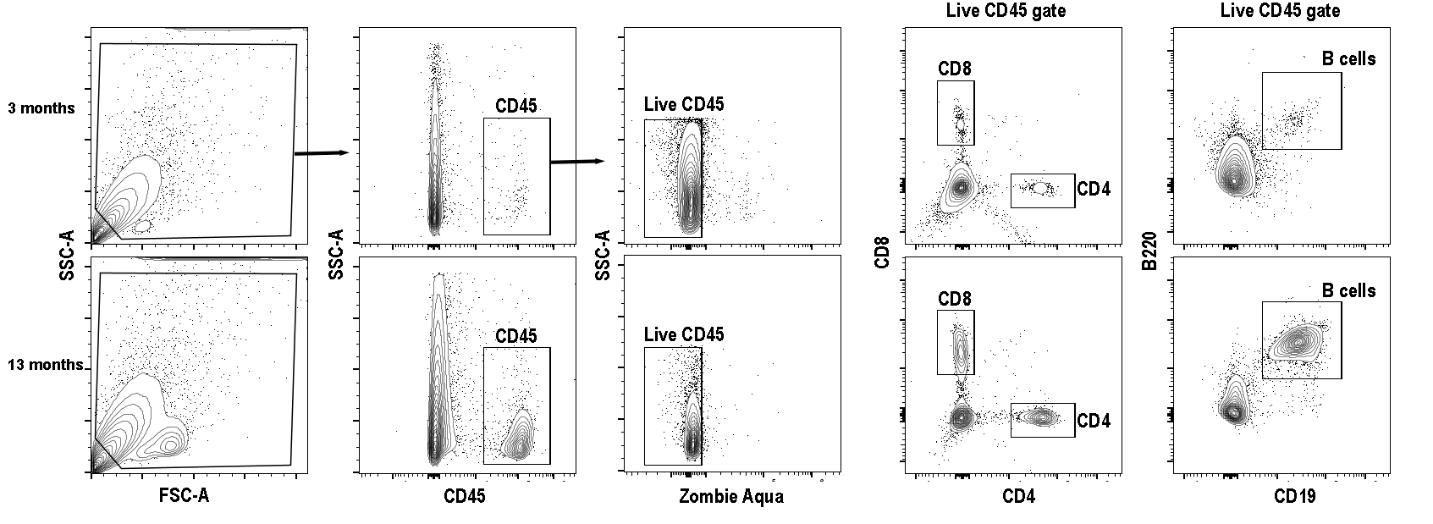

**Supplementary Figure S3B.** Analysis of immune cells infiltrating the salivary glands at different ages. Representative plots showing the gating strategy for CD45+, live, CD4 T, CD8 T, and CD19+B220+ B cells at 3- and 13- months.

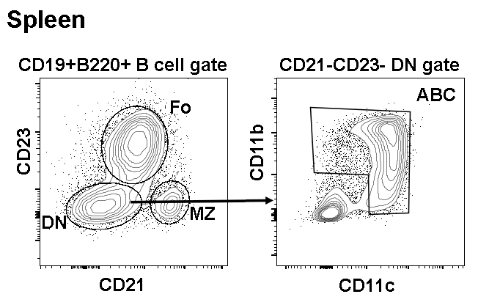

**Supplementary Figure S4.** Gating strategy showing CD23+CD21+ Follicular cells (Fo), CD21hi CD23- Marginal Zone (MZ) cells, and CD21-CD23- Double Negative (DN) cells within the B cell gate. ABCs were identified as the DN cells expressing either CD11b and/or CD11c.

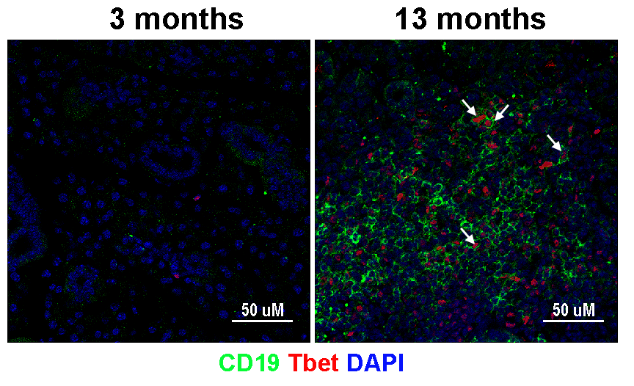

**Supplementary Figure S5.** Representative images of salivary gland sections stained with antibodies to CD19 and T-bet at 3- and 13-months showing T-bet expression within B cells within the lymphocytic focus (arrows). T-bet is also seen in other immune infiltrating cells within the focus.

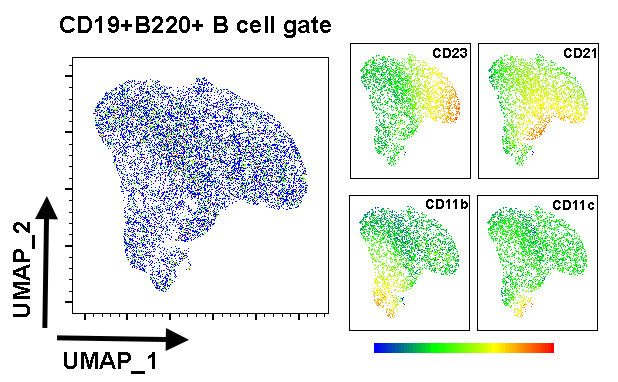

**Supplementary Figure S6.** UMAP plots with multicolor plot overlays showing CD11b, CD11c, CD21, and CD23 intensities. The plot was created by concatenating 15,000 cells in the CD19+B220+ gate from salivary glands, spleens, and livers from 8 and 13-month-old mice, n=5/age group,500 cells/mouse.

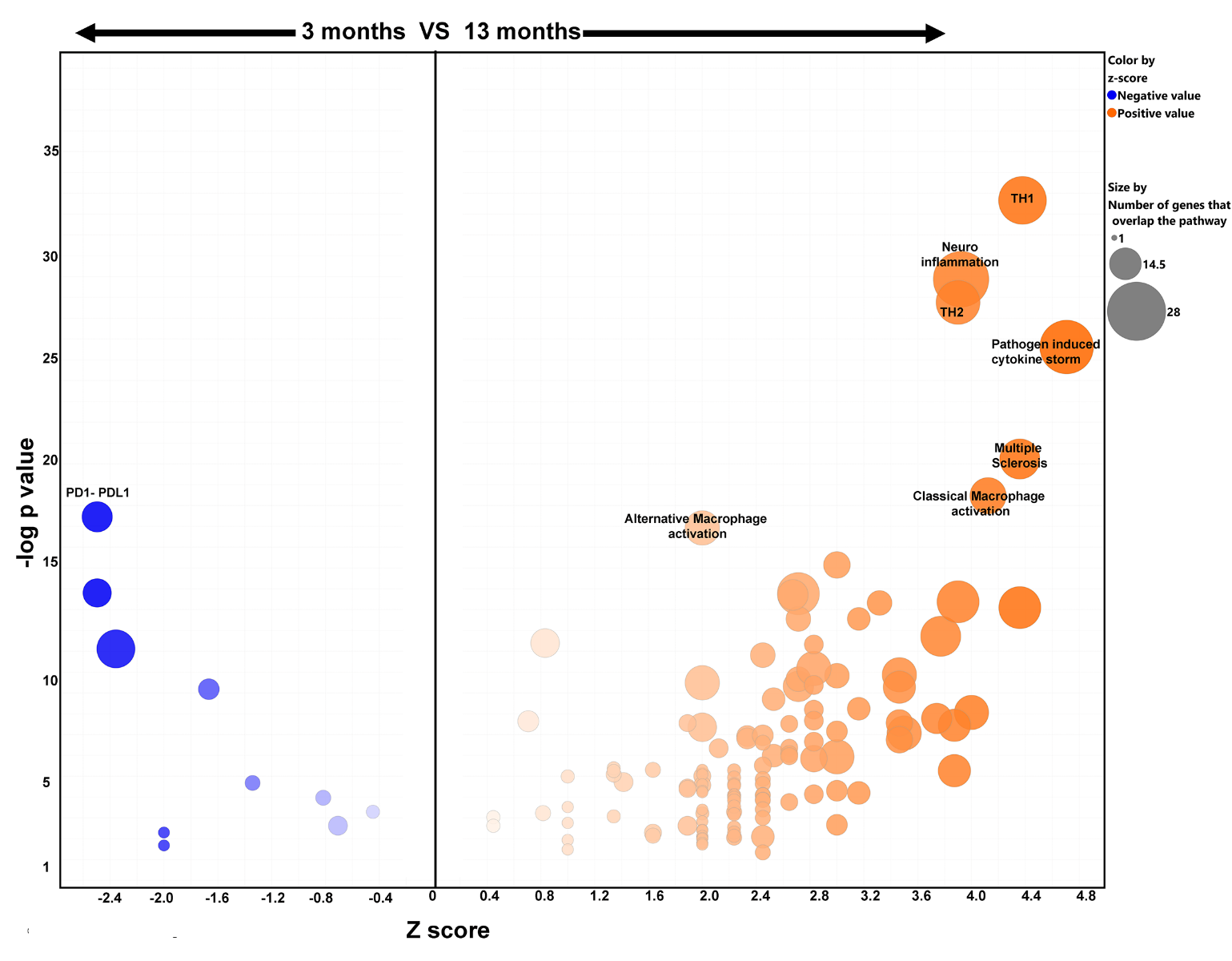

**Supplementary Figure S7.** Ingenuity pathways analysis of DE genes from salivary glands at 3- and 13-months of age showing Z scores and p values for each pathway. The top 7 signaling pathways with positive Z scores indicating enrichment at 13 months are labeled. The most significantly downregulated PD1-PDL1 pathway with a negative Z score at 13 months is shown.

**Supplementary Tables**

**Supplementary Table S1: Mouse substrains and source**

| Strain  Supplier/Source | C57BL6/J  (JAX, OMRF) | C57BL6/N  (NIA, CRL) | Total |
| --- | --- | --- | --- |
| Age (months) | **n** | **n** | **n** |
| 3 | 8 | 5 | 13 |
| 8-9 | 9 | 5 | 14 |
| 13-14 | - | 10 | 10 |
| 17-26 | 8 | 10 | 18 |

**JAX:**	The Jackson Laboratory

**NIA:**	National Institute of Aging Aged Rodent Colonies maintained at Charles River Laboratories

**CRL:**	Charles River Laboratories

| **Supplementary Table S2. List of antibodies used for immunohistochemistry (PLP fixed OCT embedded cryosections)** | | | | | |
| --- | --- | --- | --- | --- | --- |
| **Antigen** | **Fluor** | **Clone** | **Supplier** | **Catalog no** | **dilution** |
| MHC II | AF 488 | M5/114.15.2 | Biolegend | 107615 | 1:400 |
| CD3 | AF 594 | 17A2 | Biolegend | 100240 | 1:400 |
| CD4 | AF595 | GK1.5 | Biolegend | 100446 | 1:100 |
| CD8 | eFluor 660 | 53-6.7 | eBiosciences | 50-0081-80 | 1:50 |
| B220 | FITC | RA3-6B2 | eBiosciences | 11-0452-82 | 1:100 |
| CD19 | AF 647 | 6D5 | Biolegend | 115522 | 1:50 |
| T-bet |  | rabbit polyclonal | AbCam | EPR27094-16 | 1:100 |
| anti- rabbit IgG | AF 488 | donkey polyclonal | Jackson Immunoresearch | 711-546-152 | 1:100 |
| **Supplementary Table S2. List of antibodies used Flow cytometry** | | | | |  |
| **Antigen** | **Fluor** | **Clone** | **Supplier** | **Catalog no** | **Titrate as needed** |
| B220 | PE-Cy7 | RA3-6B2 | eBioscience | 25-0452-82 |  |
| CD103 | BV785 | 2E7 | Biolegend | 121439 |  |
| CD11b | PE-Cy5 | M1/70 | Biolegend | 101210 |  |
| CD11c | BV711 | N418 | Biolegend | 117349 |  |
| CD138 | BV650 | 281-2 | Biolegend | 142518 |  |
| CD19 | PE | 1D3 | BD Biosciences | 553786 |  |
| Cd19 | BUV661 | 1D3 | BD Biosciences | 612971 |  |
| CD206 | BV605 | C068C2 | Biolegend | 141721 |  |
| CD21 | APC-Fire 750 | 7E9 | Biolegend | 123433 |  |
| CD23 | PE Dazzle 594 | B3B4 | Biolegend | 101633 |  |
| CD278 | PE-Cy5 | 15F9 | BioLegend | 107708 |  |
| CD279 | PE Dazzle 594 | 29F.1A12 | BioLegend | 135227 |  |
| CD4 | BUV563 | GK1.5 | BD Biosciences | 612923 |  |
| CD44 | AlexaFluor 488 | IM7 | BioLegend | 103015 |  |
| CD45 | BUV737 | 30-F11 | BD Biosciences | 748371 |  |
| CD49a | BV421 | Ha31/8 | BD Biosciences | 740046 |  |
| CD49b | APC | DX5 | BioLegend | 108909 |  |
| CD5 | PE | 53-7.3 | ThermoFisher | 12-0051-82 |  |
| CD62L | BV786 | MEL-14 | BioLegend | 104440 |  |
| CD64 | BV421 | X54-5/7.1 | Biolegend | 139309 |  |
| CD8 | BUV395 | 53-6.7 | BD Biosciences | 563786 |  |
| F4/80 | AlexaFluor 647 | BM8 | eBioscience | 51-4801-82 |  |
| FceR1 | PE | MAR-1 | BioLegend | 134308 |  |
| GL-7 | PerCp- Cy5.5 | GL7 | BioLegend | 144609 |  |
| IgD | FITC | 11-26c.2a | Biolegend | 405704 |  |
| Ly6C | APC -Fire 810 | HK1.4 | Biolegend | 128055 |  |
| MHC II | BUV 805 | M5/114.15.2 | BD Biosciences | 748844 |  |
| NK1.1 | APC- Fire 750 | PK136 | BioLegend | 108752 |  |
| Siglec H | PE | 551 | Biolegend | 129605 |  |
| TCR beta | PE | H57-597 | BD Biosciences | 553172 |  |
| TCR gd | PE | GL3 | BioLegend | 118108 |  |
| Ter119 | PE | TER-119 | BioLegend | 116208 |  |
